# Enhancing the translational capacity of *E. coli* by resolving the codon bias

**DOI:** 10.1101/318105

**Authors:** Zoltan Lipinszki, Viktor Vernyik, Nora Farago, Tobias Sari, Laszlo G. Puskas, Frederick R. Blattner, Gyorgy Posfai, Zsuzsanna Gyorfy

## Abstract

*Escherichia coli* is a well-established, and popular host for heterologous expression of proteins. The preference in the choice of synonymous codons (codon bias), however, might differ for the host and the original source of the recombinant protein, constituting a potential bottleneck in production. Codon choice affects the efficiency of translation by a complex and poorly understood mechanism. The availability of certain tRNA species is one of the factors that may curtail the capacity of translation.

Here we provide a tRNA-overexpressing strategy that allows the resolution of the codon bias, and boosts the translational capacity of the popular host BL21(DE3) when rare codons are encountered. In BL21(DE3)-derived strain, called SixPack, copies of the genes corresponding to the six least abundant tRNA species have been assembled in a synthetic fragment and inserted into a ribosomal RNA operon. This arrangement, while not interfering with the growth properties of the new strain, allows dynamic control of the transcription of the extra tRNA genes, providing significantly elevated levels of the rare tRNAs in exponential growth phase.

Results from expression assays of a panel of heterologous proteins of diverse origin and codon composition showed that the performance of SixPack surpassed that of the parental BL21(DE3) or a related strain equipped with a rare tRNA-expressing plasmid.

**Importance:** Codon composition not fitting the codon bias of the expression host frequently compromises the efficient production of foreign proteins in *E. coli*. Various attempts to remedy the problem (codon optimization by gene synthesis, expression of rare tRNAs from a plasmid) proved to be unsatisfying. Our new approach, adjusting the tRNA pool by co-expressing extra copies of rare tRNA genes with ribosomal RNA genes, does not affect normal cell physiology, and seems to be a superior solution in terms of simplicity, cost, and yield.

## Introduction

*E. coli* is by far the most widely used host organism for biopharmaceutical heterologous production of recombinant proteins. This expression platform is favored for its simplicity, speed, and low cost. The codon bias discrepancy, however, can seriously hinder protein expression in *E. coli* (1–5). Choice in the usage of synonymous codons can be different in various organisms, and this bias has been shown to correlate with the relative and absolute quantities of individual tRNAs (6,7). Heterologous expression of a protein with a high ratio of codons occurring infrequently in *E. coli* might deplete the corresponding tRNA species, leading to translational frameshifting, codon skipping, misincorporations, and protein truncations (8). Ultimately, the codon bias seriously limits the use of *E. coli* as an expression platform.

To overcome this problem, codon optimization or rare tRNA overexpression strategies have been applied, with limited success. On one hand, synthesis of the recombinant protein encoding gene with an *E. coli* codon preference is labor-intensive and expensive. Moreover, replacing rare codons with frequent ones does not necessarily lead to increased yield in protein synthesis. Codon choice might affect expression, solubility, and folding of a protein (9–12); and rare codons can, paradoxically, enhance the translation of a gene via reducing the mRNA secondary structure (7,13,14). Additionally, codon optimization by gene synthesis is not feasible when testing gene libraries. On the other hand, attempts to overexpress rare tRNA species by cloning extra copies of the corresponding tRNA encoding genes into a plasmid have their drawbacks as well. In such commercially available hosts, maintenance of the tRNA-expressing plasmid requires the addition of extra antibiotics (usually the protein synthesis inhibitor chloramphenicol); moreover, the recombinant protein-encoding expression plasmid must belong to a different complementation group. Using two antibiotics and/or permanently altering the balance of the various tRNA species can have a fitness cost on the host, eventually resulting in a low success rate in applications (10,15).

We sought to resolve the codon bias problem by expressing rare tRNAs in a more dynamic fashion. We hypothesized that inserting extra copies of the relevant tRNA genes into a ribosomal RNA (*rrn*) operon would harmonize their expression with the translational activity and would provide enhanced levels of rare tRNAs according to the actual needs. *rrn* operons have key roles in bacterial physiology and economy. Synthesis of rRNA quickly reacts to environmental conditions, determining ribosome availability via regulating the expression of ribosomal proteins by a translational feedback mechanism (16). In *E. coli*, there are seven nearly identical copies of rRNA operons. Synthesis of rRNA is driven by the strongest promoters found in the genome (17), and the rate of transcription can change more than an order of magnitude between poor and rich nutrient conditions (18). While most of the tRNA genes are scattered around the chromosome, all rRNA operons carry certain (but not any of the rare) tRNA genes as well, co-transcribed with the rRNA genes.

We inserted extra copies of the six tRNA genes (*argX*, *glyT*, *leuW*, *proL*, *argU,* and *ileX*) corresponding to the minor codons of *E. coli* (CGG, GGA, CUA, CCC, AGA/AGG, and AUA, respectively) into one of the ribosomal RNA operons (*rrnD*) of BL21(DE3), the widely used *E. coli* expression host. We demonstrated that expression of these tRNA genes varies with growth rate and shows a marked increase compared to the unmodified host. By testing the expression of a panel of recombinant proteins with a high ratio of rare codons in their genes, we showed that, in most cases, the modified strain named SixPack performs better and shows significantly enhanced expression of the heterologous proteins in classical IPTG-induction as well as in an auto-induction system. Moreover, compared to the commercially available strain Rosetta2(DE3)pLysS (Merck), which carries a similar array of extra tRNA genes on a plasmid (pLysSRARE2; Merck), SixPack proves to be a superior expression platform.

## Results

### Design and genomic insertion of extra copies of tRNA genes

Hypothetically, cloning extra copies of the genes corresponding to rare tRNAs into a *rrn* operon allows their controlled expression, as the activity of the operon is tightly regulated by nutrient availability and other physiological conditions (19). This might ensure that expression of the rare tRNAs does not cause a permanent extra burden for the cell.

The genes encoding for the six least abundant tRNA species were combined in a single, synthetic DNA fragment. The original copies of these genes are either single genes whose expression is controlled by their own promoter (*proL*, *argU,* or *ileX*), or are parts of polycistronic operons (*argX*, *glyT,* or *leuW*) (Fig. S1). Since the process of maturation of the primary transcripts of tRNA genes is not thoroughly understood, the nucleotide sequences surrounding the genes in the synthetic fragment were designed to mimic the natural tRNA operons. The intergenic and flanking regions of the *argX-glyT-leuW-proL* segment were essentially identical to those of the natural *argX-hisR-leuT-proM* polycistronic operon. For *argU* and *ileX*, exact copies of the genes and their flanking sequences were fused to the 3’ end of the segment (Fig. S2).

For the genomic insertion site *rrnD*, one of the seven *rrn* operons was selected. Although the structure, sequence, and activity of the *rrn* operons of *E. coli* are very similar, the 3’ end of *rrnD* (containing *trhV* and *rrfF*) is unique, allowing specific targeting of the locus (17).

The 1207-base synthetic operon carrying the six tRNA genes was engineered into the 3’ region of *rrnD* (Fig. 1) by a multistep process using λRED recombineering, followed by elimination of the markers via CRISPR-Cas9-stimulated homologous recombination (Fig. S3), resulting in strain SixPack.

**Figure 1.**
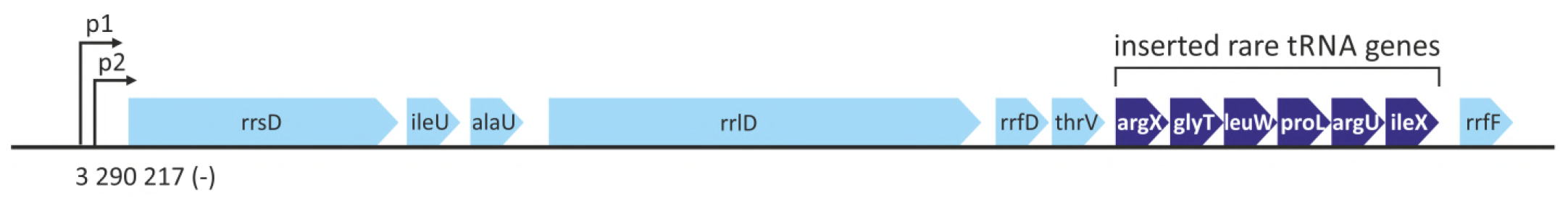
Schematic picture of *rrnD* showing the six extra tRNA genes (dark blue) inserted in the 3’ end region between *thrV* and *rrfF*.

### Growth properties of SixPack and control strains

Apparently, insertion of the extra tRNA genes into *rrnD* did not disturb the homeostasis of the cell. The growth parameters of SixPack were identical to those of the parental BL21(DE3) in both LB and AIM (auto-induction medium). Noticeably, the commercially available Rosetta2(DE3)pLysS strain had a longer lag phase, lower growth rate, and lower maximum OD in both media, presumably owing to the burden of maintaining the plasmid and the presence of the antibiotic chloramphenicol in the media (Fig. 2 and Table S1).

**Figure 2.**
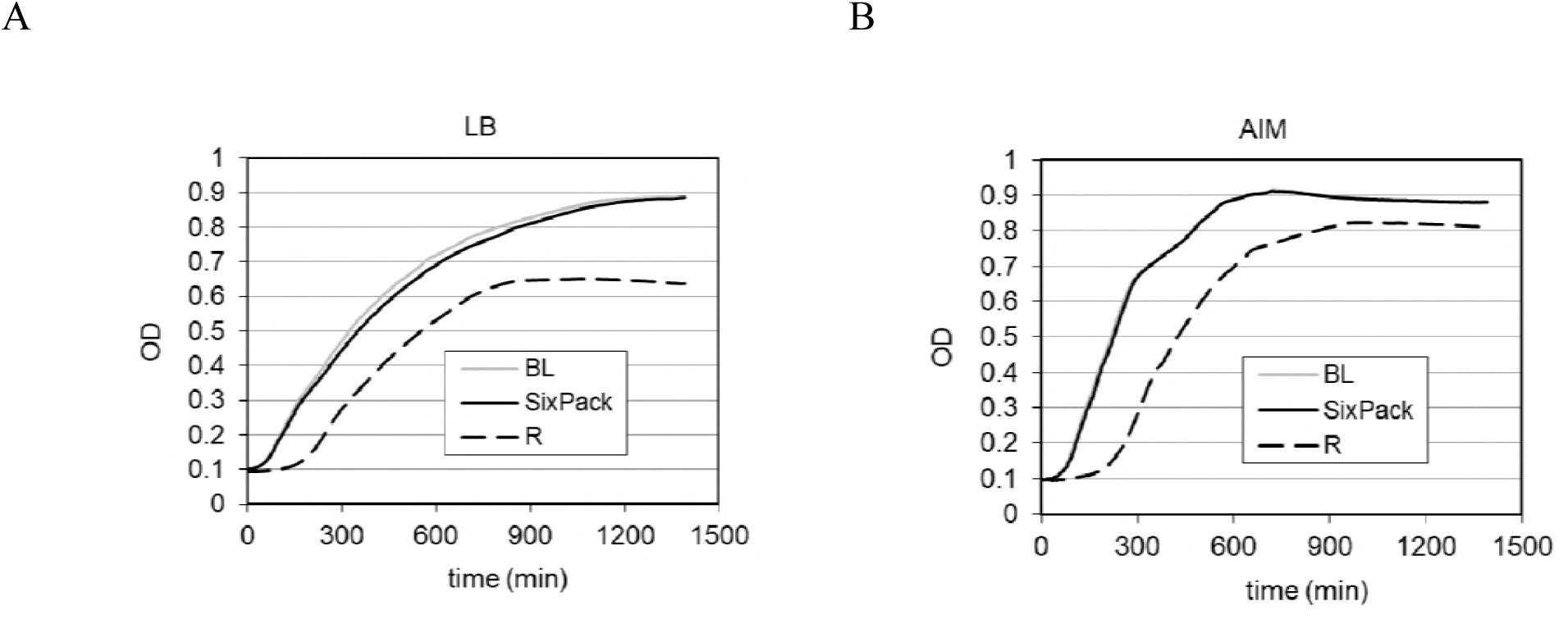
Growth curves of SixPack and control strains BL21(DE3) (marked BL) and Rosetta2(DE3)pLysS (marked R) in LB **(A)** and in AIM **(B)**. (Averages of three independent experiments.)

### Elevated expression of rare tRNAs

Expression of the six rare tRNA genes was measured by qRT-PCR on first-strand cDNA generated from total RNA isolated from cultures of BL21(DE3) and SixPack in the exponential phase (OD_600_ = 0.45) and in the early stationary phase (OD_600_ = 4.5). Primers were designed to amplify both the unprocessed and the matured forms of the various tRNA species. In the case of SixPack, the measured rare tRNAs represented transcripts originating both from their original genomic copies and from the newly inserted fragment in *rrnD*. The rare tRNA ratios were normalized with the ratios of three control (abundant) tRNA species.

Compared to parental BL21(DE3), SixPack showed elevated expression of the six rare tRNAs. The difference was more pronounced in the exponential phase (1.4- to 3.8-fold) than in the stationary phase (0.8- to 3.0-fold) (Fig. 3).

**Figure 3.**
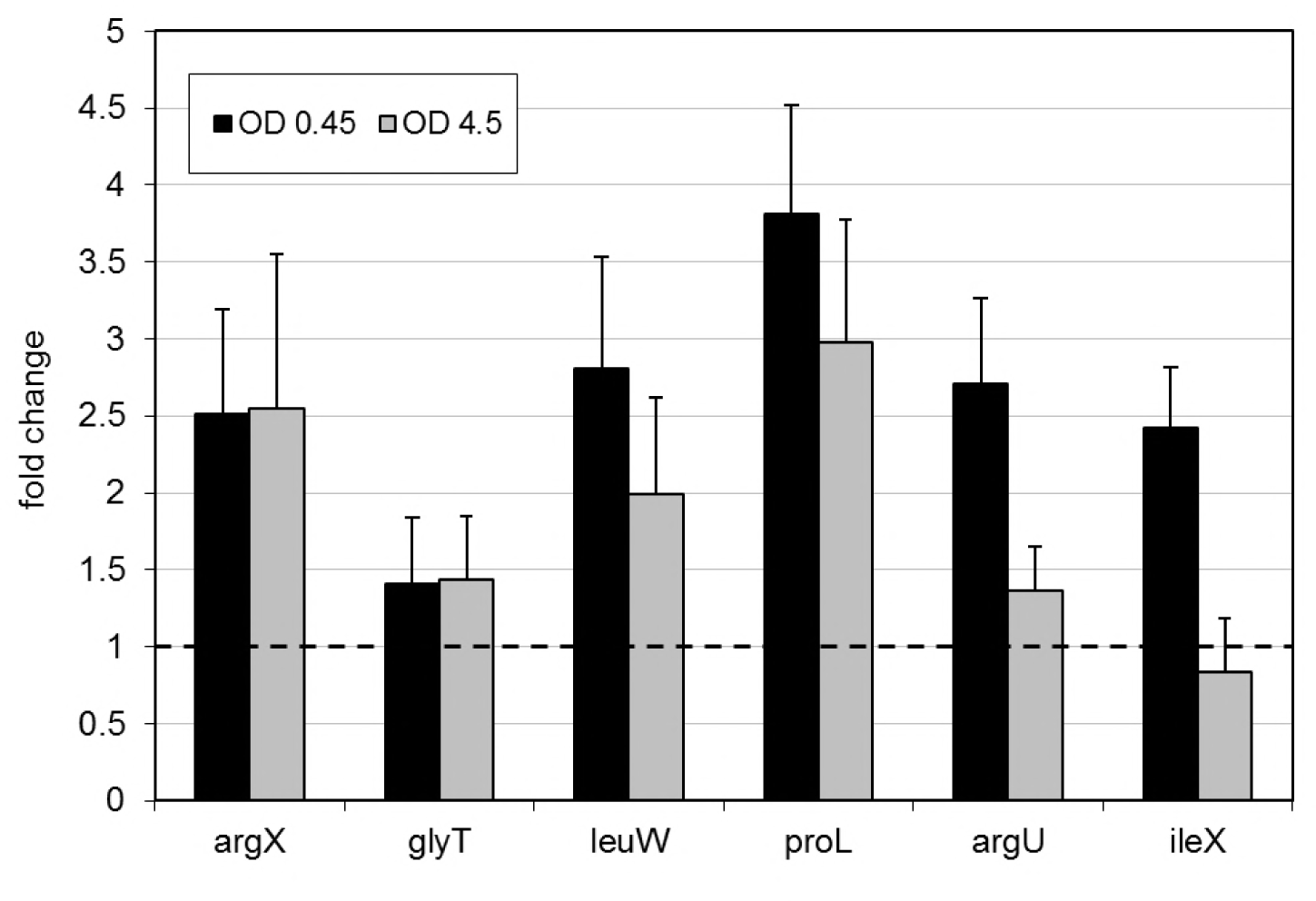
Ratios of the different tRNA species between SixPack and BL21(DE3) in the exponential growth phase (black columns) and in the early stationary phase (gray columns). (Averages of two independent experiments, each comprising three technical repetitions.)

Despite being on the same cistron and transcribed together, increases in the expression levels of the extra tRNA genes were diverse (*proL* showing the highest and *glyT* the lowest increase). This might be due to several factors, including tRNA processing, stability, regulation, and diverse expression levels of the original genomic copies.

### Individual impact of elevated levels of the rare tRNAs on protein expression

The functional effect of the increased levels of rare tRNAs was first tested separately for each tRNA species. To magnify the effect of rare tRNAs, we constructed IPTG-inducible, pETDuet-1-based (Novagen) test plasmids expressing modified versions of the GFP gene under the regulation of the T7 promoter. GFP genes carrying runs of rare codons corresponded to each of the six extra tRNA genes (one GFP version corresponding to *proL*, *ileX*, *argX*, *glyT,* and l*euW* each, and two GFP versions for *argU*, as it relates to both AGA and AGG codons). The tandem rare codons (3 or 5 copies) were inserted into the GFP gene immediately after the beginning ATG codon. Expressions of the modified GFPs were detected by fluorescence measurements and visualized by denaturing polyacrylamide gel electrophoresis (SDS-PAGE).

IPTG- (or lactose-) induced expression of the GFP and its variants with rare codons had a negative but diverse impact on the growth of the strains. In LB medium, SixPack and BL21(DE3) usually showed similar growth patterns, and in most cases displayed higher growth parameters than Rosetta2(DE3)pLysS. In AIM, peculiarly, Rosetta2(DE3)pLysS reached the highest maximal optical density when GFP variants with rare codons were expressed (Fig. S4 and S5).

Total fluorescence measurements of GFP production, however, showed a clear advantage of SixPack in both LB (Fig. 4) and AIM (Fig. S6). In both media, SixPack showed higher expression of the modified GFPs than the parental BL21(DE3) did. In one exception, there was no difference (3xGGA-GFP; GGA codon relates to *glyT*) (Fig. S6H). Compared to Rosetta2(DE3)pLysS, SixPack showed higher expression in all cases.

**Figure 4.**
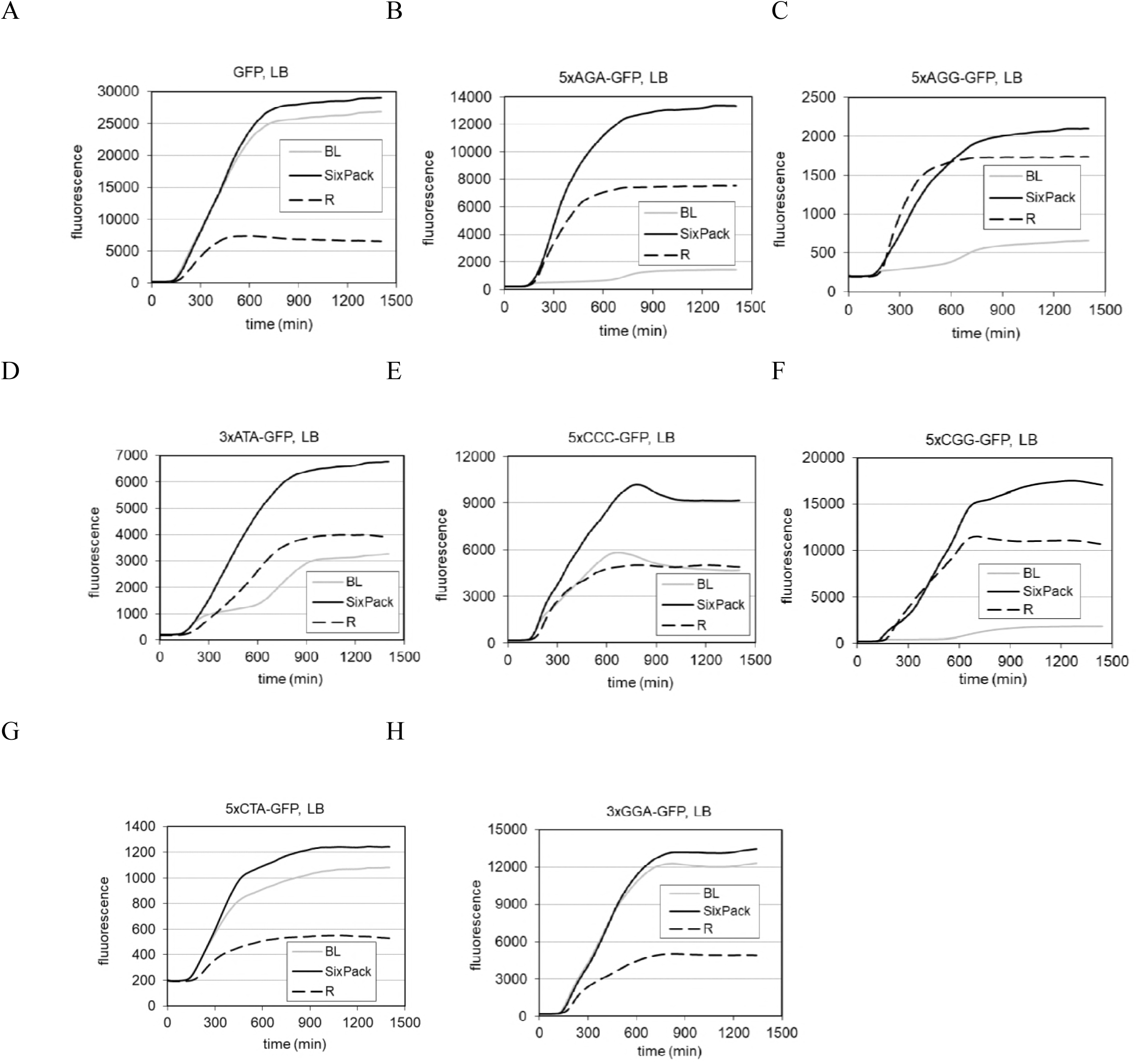
Expression of GFP (**A**) and modified GFP versions (**B-H**) in BL21(DE3) (marked BL), SixPack, and Rosetta2(DE3)pLysS (marked R) in LB, monitored by fluorescence measurements. (The curves represent the averages of four independent experiments.)

Results of the fluorescence measurements were further supported by protein gel assays. Equal amounts of proteins obtained from crude extracts of cultures collected after a 10-h incubation were analyzed on Coomassie Brilliant Blue-stained SDS-PAGE gels. Differences in the amounts of modified GFPs in the three strains were in clear correlation with the fluorescence data (Fig. S7).

### Heterologous protein expression performance of SixPack and control strains

The usefulness of SixPack was further tested by expressing a panel of eight protein-coding genes of diverse length and rare codon content (Table S2), selected from various species, including *Drosophila melanogaster, Pyrococcus furiosus*, *Saccharomyces cerevisiae*, *Streptococcus pyogenes,* and human papillomavirus (HPV). The genes were cloned into pETDuet-1 and expressed in SixPack and control strains under two different conditions (LB and AIM).

The growth curves of IPTG- (or lactose-) induced cultures seemed to be quite diverse in LB (Fig. S8), but rather uniform in AIM (Fig. S9). Production of the specific proteins was visualized by using Coomassie-stained gels and Western blotting (Fig. 5). In most cases, SixPack proved to be the best producer. This was especially evident when the ratio of rare codons was higher than 8% (and the ratio of tandems was also high), as in the cases of HumdCas9 (dCas9 optimized to human codon preference), HumdCas9-GFP fusion, and *Pfu* DNA polymerase. In the case of EFT1, the ratio was only 4.3% (35 AGA and 2 CTA codons); but SixPack still produced the highest amount of this protein. It is conceivable that the translation efficiency is influenced not only by the total amount of rare codons, but also by the occurrence of tandem situations, the location of rare codons in the gene (e.g., close or far from the start site), and the ratios of the different rare codon species (3). In other cases (Asl, dCas9, Flfl, and HPV16 L2), when the rare codon ratio was below 8%, production by SixPack was similar to that of BL21(DE3) in LB and similar to or better than that in AIM. In all cases, the performance of Rosetta2(DE3)pLysS lagged behind that of SixPack (especially evidently in AIM) or was similar at most (Fig. 6).

**Figure 5.**
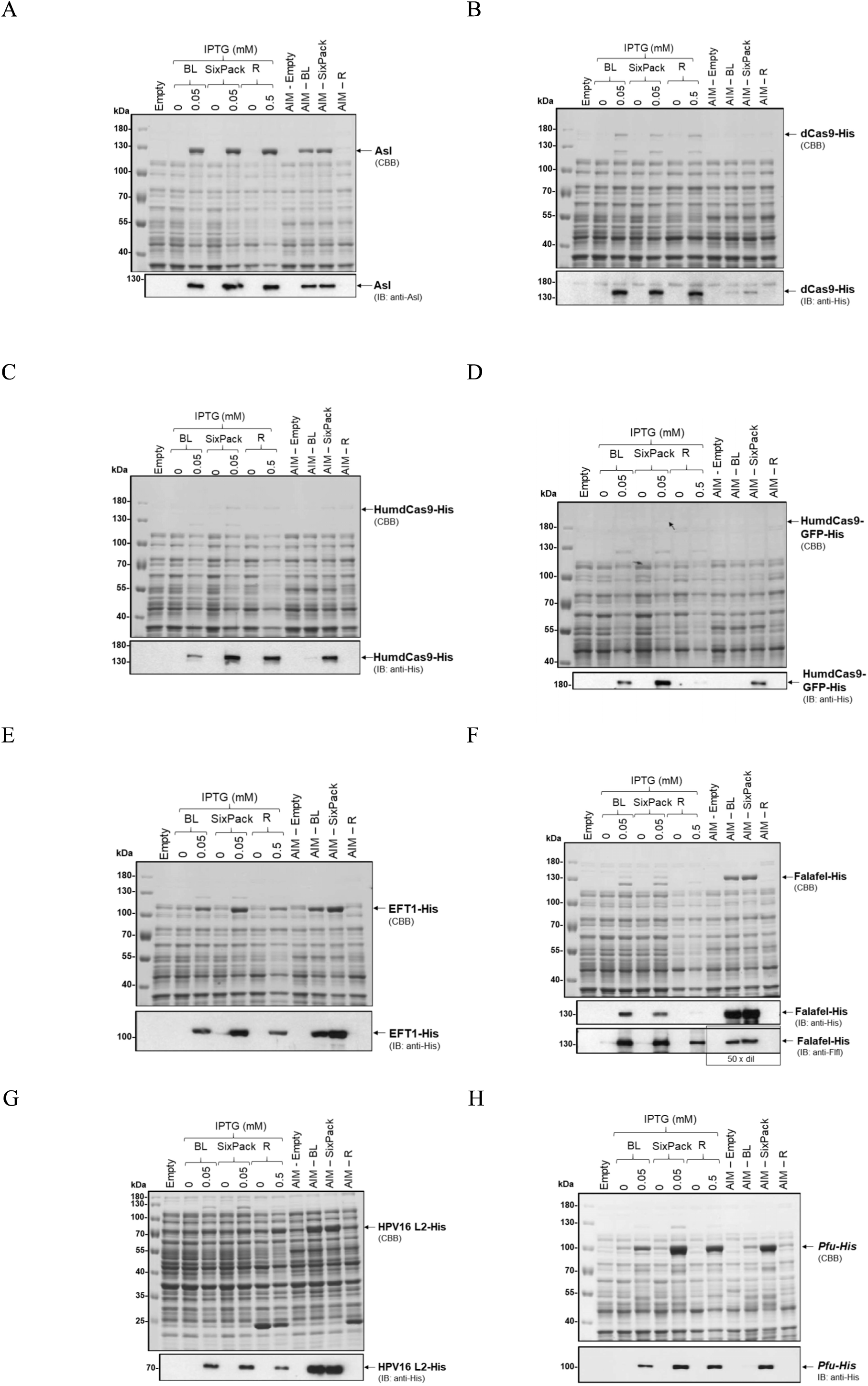
Production of eight heterologous proteins (**A-H** for Asl, dCas9, HumdCas9, HumdCas9-GFP fusion, EFT1, Flfl, HPV16 L2, and *Pfu*, respectively) detected on protein gels by Coomassie Brilliant Blue staining and Western blotting. Proteins were extracted from BL21(DE3) (marked BL), SixPack and Rosetta2(DE3)pLysS (marked R) after 10 h growth in LB or AIM.

**Figure 6.**
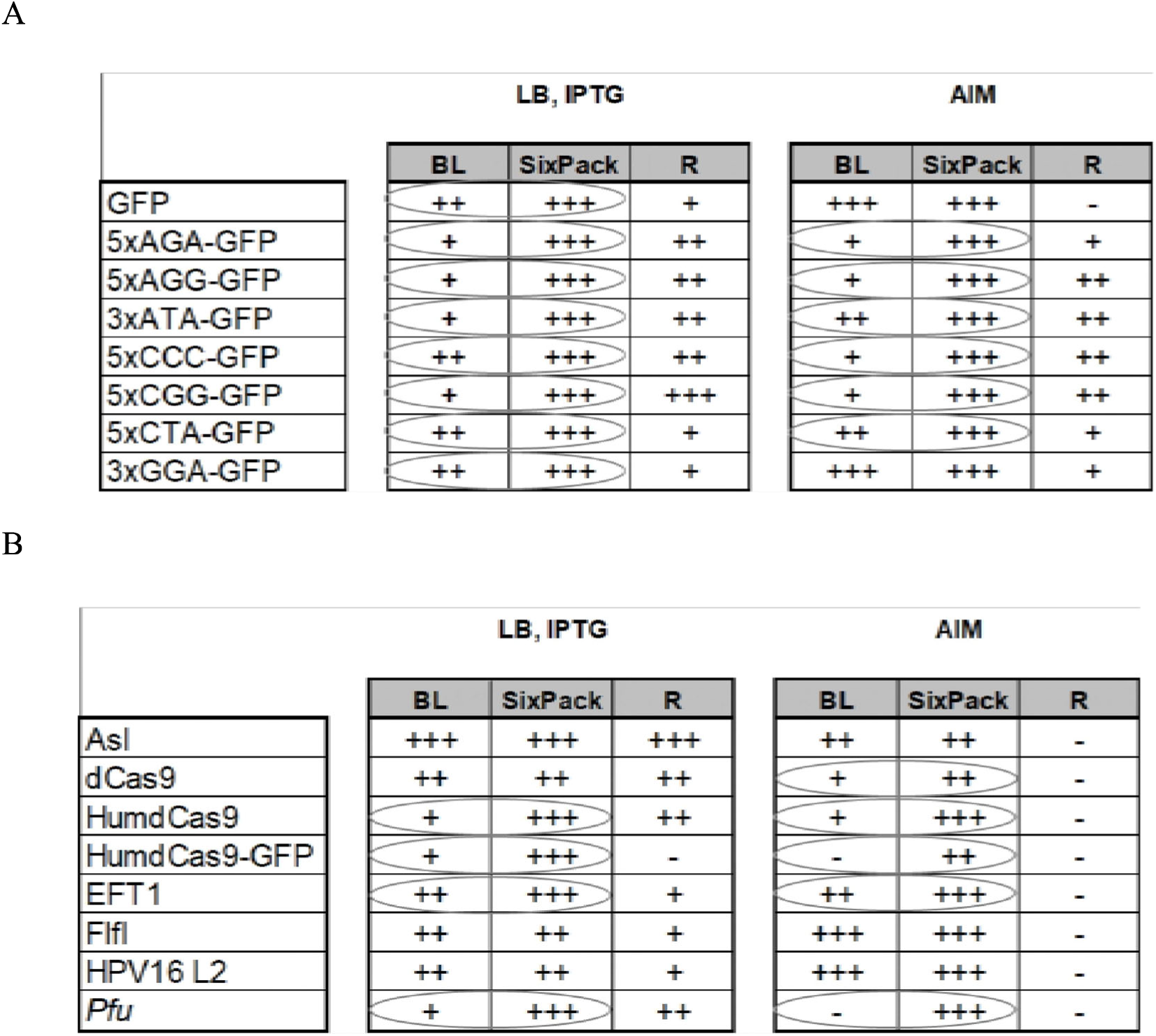
Summary of heterologous protein production of BL21(DE3) (marked BL), SixPack, and Rosetta2(DE3)pLysS (marked R) measured in LB and AIM. The “+” and “-“ marks indicate the relative protein production of the strains. Table **A** is based on the maximal fluorescence values of GFP and GFP variants detected in the first 24 h after induction. Table **B** is based on results of Western blots - validated by densitometry (data not shown). Circles label the cases when SixPack performed better than the parental strain BL21(DE3).

## Discussion

Depletion of rare tRNA species evoked by the expression of heterologous proteins can compromise product yield and translational fidelity. Several attempts, both genetic (overexpression of rare tRNA genes from a plasmid, or codon optimization of the heterologous gene by synthesis) and non-genetic ones (optimizing the growth conditions, or using *in vitro* protein synthesis systems) have been applied with varying degrees of success (20–23). We offer here a solution that resolves codon bias discrepancies with a high success rate, does not require additional antibiotics, and performs better or as well as the widely used BL21(DE3) *E. coli* strain.

The novelty of the approach lies in placing the extra copies of rare tRNA genes into a ribosomal operon of the host chromosome. This arrangement offers the following benefits related to the rare tRNA expression: (i) lack of physiological burden associated with plasmid replication and maintenance; (ii) genetic stability without the use of an antibiotic; and (iii) dynamic changes in the levels of rare tRNAs, paralleling the actual translational activity. Consequently, elevated levels of rare tRNAs are achieved with minimal interference with normal host physiology.

A ribosomal operon is a natural choice as a location for inserting tRNA genes. While most tRNA genes are scattered elsewhere in the genome, each of the *rrn* operons carries at least one tRNA gene (typically coding for abundant tRNAs). Among the seven *rrn* operons, *rrnD* is particularly suited for insertion of the extra genes, as it carries unique regions near its 3’ end, allowing easy targeting. In this vein, a synthetic segment carrying the genes of the six least abundant tRNAs of *E. coli* (*argX*, *glyT*, *leuW*, *proL*, *argU*, *ileX*) was cloned into *rrnD*, downstream of its own tRNA gene *thrV*.

As anticipated, increased amounts of the rare tRNAs, corresponding to the inserted genes, were detected by qRT-PCR. Moreover, their levels showed a growth phase-dependent adjustment (higher in exponential phase, lower in stationary phase). However, when compared to the levels detected in the parental host BL21(DE3), the rate of increase in the exponential phase was only modest (1.4- to 3.8-fold), especially when considering the wide dynamic range of *rrn* transcripts across the growth cycle. An interplay of several factors, including co-factors of tRNA maturation, tRNA stability, and feedback regulation of the genes at the original locations, might mitigate the increase.

Although the increase in the abundance of the rare tRNAs did not interfere with cell growth, it had a significant, positive effect on heterologous protein expression. Individually, with the exception of *glyT* in AIM, all the genes caused increased expression (up to 12-fold) of specific GFP variants carrying tandem copies of the corresponding rare codon. The combined effect of the six extra genes was then clearly demonstrated by expressing a panel of proteins of various origin and codon bias. Expression in SixPack (amount of target protein per cell mass or per culture volume) was at least as good as in any of the control strains and in fact, was better in most cases (in some cases even 20-fold). In general, with the increasing number of rare codons and their tandem copies, the advantage of SixPack became increasingly evident. SixPack compared especially favorably to control strains when applying AIM, which is a popular medium for biotechnological production of proteins in large volumes (i.e., in bioreactors) (24).

Insertion of the six rare tRNA genes into the genome of BL21(DE3) did not result in changes in its basic characteristics like morphology, growth properties, or transformation efficiency. The beneficial features (no need for extra plasmid and antibiotics, no need for codon optimization, simplicity when expressing gene libraries, high success rate when expressing proteins with different codon bias) render SixPack a useful, and likely superior, alternative host for expression of heterologous proteins.

## Materials and Methods

### Medium

In all experiments involving bacterial culture growth, standard LB or AIM (auto-induction medium; LB broth base including trace elements; Formedium LTD, England) were used. Antibiotics were used in the following concentrations: 100 μg/ml ampicillin (Ap) and 24 μg/ml chloramphenicol (Cam).

### *E. coli* strains

BL21(DE3) was used as the parental strain. It expresses T7 RNA polymerase from the λDE3 lysogen inserted into the genome under the regulation of the *lacUV5* promoter. This polymerase can drive the expression of the genes of interest via the T7 promoter (25). Rosetta2(DE3)pLysS is a derivative of BL21(DE3). In this strain, pLysSRARE2 plasmid encodes seven genes of rare tRNAs (*argU*, *argX*, *argW*, *glyT*, *leuW*, *ileX,* and *proL*, recognizing the AGA/AGG, CGG, AGG, GGA, CUA, AUA, and CCC codons, respectively) under the control of their own promoter. pLysSRARE2 also contains chloramphenicol resistance- and lysosyme-encoding genes. Due to constitutive lysosyme expression, the T7 polymerase-driven expression of the genes of interest is lower in Rosetta2(DE3)pLysS than in BL21(DE3) or SixPack (26). To create the SixPack strain, BL21(DE3) was modified by inserting extra copies of six rare tRNA genes (*argU*, *argX*, *glyT*, *leuW*, *ileX,* and *proL*) into its *rrnD* operon (Fig. 1). These tRNAs recognize the same rare codons as the tRNAs expressed from the pLysSRARE2 plasmid (AGG is recognized by *argX* and *argW* in Rosetta2(DE3)pLysS, but only by *argX* in SixPack).

### Genomic construct

The rare tRNA genes were cloned into pSG76A (replicating by R6K ori) in three pieces. The first part (synthesized by Thermo Fisher Scientific GENEART GmbH; Germany) contained the *argX*, *glyT*, *leuW,* and *proL* tRNA genes. The structure of this DNA fragment was based on the polycistronic operon of *E. coli* coding for *argX*, *hisR*, *leuT,* and *proM* tRNA genes; we replaced the *hisR*, *leuT,* and *proM* genes with *glyT*, *leuW,* and *proL* rare tRNA genes, but kept *argX* and the original intergenic regions. The second and third pieces coding for *argU* and *ileX*, respectively, were directly amplified from the *E. coli* genome with their own intergenic regions (Fig. S1 and S2).

The plasmid carrying the six tRNA genes was linearized by PCR using overhanging primers containing homologous regions (50- or 100-nt) with the target site (unique sequence of the 3’ end of *rrnD*), and transformed into BL21(DE3) in the presence of λRED recombinase (27). After successful recombination into the genome, the CRISPR-Cas9 system was used to introduce a double-strand brake in the ampicillin resistance gene on the insert (28). The cells’ own RecA system repaired the cleavage by homologous recombination using the 47-nt box on the plasmid homologous to the adjacent *rrnD* sequence downstream from the site of insertion (29) (Fig. S3 and S10).

### Expression plasmids

The coding sequences of different test proteins were cloned into pETDuet-1 (Novagen). A list of these genes and the cloning sites is given in Table S3.
The GFP-encoding gene was amplified from pCA24N (the cloning plasmid of the ASKA collection; 30), and subcloned into pETDuet-1. GFP gene variants containing runs of rare codons were created by PCR using pETDUET/GFP as a template.
The coding DNA sequence of asterless (Asl) was subcloned from the GH02902 Drosophila Gold cDNA clone into pETDuet-1 following standard procedures.
Cloning of Flf1 into pETDuet-1 has been described elsewhere (31).
dCas9 was cloned from the pdCas9 plasmid # 46569 from Addgene (32).
The HumdCas9 gene was amplified from pMLM3705 plasmid # 47754 from Addgene (33).
pETDUET/HumdCas9-GFP expresses HumdCas9 and GFP as a fusion protein. EFT1, the gene of elongation factor 2, was amplified from genomic DNA of *Saccharomyces cerevisiae* strain EMY 74.7 (34) and was a kind gift from Dr. Tamas Feher.
The HPV16 L2 gene (subcloned in Addgene plasmid # 72473) was a kind gift from Dr. Vilmos Tubak.
The gene of *Pfu* DNA polymerase was amplified from *Pyrococcus furiosus* genomic DNA (DSM 3638) which was a kind gift from Dr. Vilmos Tubak.
BL21(DE3) harbouring an empty pETDuet-1 was used as a control.

### Quantitative real-time PCR (qRT-PCR)

RNA was isolated from exponential phase (OD_600_ = 0.45) and early stationary phase (OD_600_ = 4.5) cultures grown in LB, using the E.Z.N.A. Bacterial RNA Kit (VWR, USA). To obtain cDNA, 1 μg of RNA was then reverse transcribed using the High-Capacity cDNA Reverse Transcription Kit (Thermo Fisher Scientific) in a final volume of 10 μl by using 1 pmole of the reverse primers (Table S4). After dilution with 20 μl of water, 1 μl of the diluted reaction mixture was used as a template in qRT-PCR with 10 μl of qPCRBIO SyGreen Mix (PCR Biosystems) and 1 μl of gene-specific primer mix, according to the following protocol: 10 min at 95°C followed by 40 cycles of 95°C for 25 sec, 60°C for 25 sec, and 72°C for 15 sec. qRT-PCR was performed in a LightCycler^®^ Nano Real-Time PCR System (Roche Diagnostics GmbH; Mannheim, Germany). Transcripts of *proK*, *alaU,* and *gltW* genes were used as house-keeping tRNA probes.

### Measuring and calculating growth parameters

To measure growth parameters, a Synergy 2 automated microplate reader machine (BioTek, USA) was used. Aliquots of 1μl each of BL21(DE3), SixPack, and Rosetta2(DE3)pLysS overnight starter cultures were transferred into 100 μl fresh LB or AIM medium in dedicated 96-well plates. Rosetta2(DE3)pLysS cultures were supplemented with Cam. Absorbance at 600 nm was measured every 5 min for 24 h at 37°C with continuous shaking. Growth parameters (length of the lag phase and the doubling time) were calculated by using previously described methods (35).

### Fluorescence measurements

Starter cultures of BL21(DE3), SixPack, and Rosetta2(DE3)pLysS strains expressing GFP or GFP variants were grown overnight in LB in a Synergy 2 automated microplate reader machine (BioTek, USA). Starter cultures (1 μl each) were re-inoculated into 100 μl fresh LB or AIM media supplemented with Ap and isopropyl β-D-1-thiogalactopyranoside (IPTG) (plus Cam, in the case of Rosetta2(DE3)pLysS). IPTG was used in concentrations optimized for highest induction (0.05 mM for BL21(DE3) and SixPack, 0.5 mM for Rosetta2(DE3)pLysS). Measurements were made every 5 min.

### Protein expression and whole cell lysate preparation

Protein expressions in BL21(DE3) and SixPack were induced with 0.05 mM IPTG, and those in Rosetta2(DE3)pLysS were induced with 0.5 mM IPTG in 3ml of LB or AIM (supplemented with antibiotics) for 10 h. Bacteria were harvested by centrifugation, resuspended in 150 μl/absorption unit (at 600 nm) of phosphate-buffered saline, pH 7.4, supplemented with 1 U/ml BenzonaseNuclease (Merck) and 2 mM MgCl_2_, kept on ice for 10 min, and then mixed with 4x Laemmli sample buffer and boiled for 5 min.

### SDS-PAGE and Western blotting

Equal amounts of protein samples were run on 7% or 10% SDS-PAGE gels. Gels were fixed for 10 min in 10% acetic acid, stained for 15 min with Coomassie Brilliant Blue (0.1% CBB in 50% methanol and 10% acetic acid), and differentiated overnight in 7% acetic acid and 10% methanol. For immunoblotting, proteins were blotted onto a nitrocellulose membrane (GE Healthcare) and probed with anti-His mouse monoclonal (Thermo Fisher Scientific, #MA1-21315; dilution 1:3-5000), anti-Flfl rat polyclonal (31) (dilution 1: 15,000), or anti-Asl rabbit polyclonal (36) (dilution 1: 20,000) antibodies following standard procedures. Stained gels and X-ray films were scanned at 600 dpi resolution for image processing.

## Acknowledgements

The authors would like to thank Dr. Vilmos Tubak for the *Pirococcus furiosus* genomic DNA and the HPV16 L2-expressing plasmid, and Dr. Tamas Feher for the EFT1-coding plasmid.

## Supplementary figure legends

Figure S1. Schematic picture of the *E. coli* BL21(DE3) genome showing the location and orientation of tRNA encoding genes (based on the BL21(DE3) complete genome sequence, CP001509.3). Circles label the rare tRNA species genes whose copies were inserted into *rrnD*.

Figure S2. Nucleotide sequence of the insert containing the six rare tRNA species genes (bold). Three incidental point mutations are underlined.

Figure S3. Main steps in the procedure for creating SixPack. The six tRNA genes and the 47-nt segment (filled orange box) homologous to the 3’ end of *rrnD* were cloned into pSG76A (**I**). The plasmid was linearized by PCR using primers bearing overhanging regions (brown, orange lines) homologous to *rrnD* (brown, orange open boxes) (**II**) and recombined into the genome via the homologous overhangs (**III**). Plasmid sequences were then eliminated by CRISPR-Cas9 cleavage-stimulated homologous recombination between the 47-nt fragment (filled orange box) and its downstream homologue (orange open box) (**IV**).

Figure S4. Growth curves of the strains expressing GFP (**A**) and different GFP variants (**B-H**) in LB medium. (The curves represent the averages of four independent experiments.)

Figure S5. Growth curves of the strains expressing GFP (**A**) and different GFP variants (**B-H**) in AIM. (The curves represent the averages of four independent experiments.)

Figure S6. Expression of GFP (**A**) and GFP variants (**B-H**) in BL21(DE3) (marked BL), SixPack, and Rosetta2(DE3)pLysS (marked R) in AIM monitored by fluorescence measurements. (The curves represent the averages of four independent experiments.)

Figure S7. Production of GFP (**A**) and GFP variants (**B-H**) after 10 h growth in LB or AIM visualized on protein gels labeled by Coomassie Brilliant Blue staining.

Figure S8. Growth curves of the strains expressing various heterologous proteins in LB. (The curves represent the averages of three independent experiments.)

Figure S9. Growth curves of the strains expressing various heterologous proteins in AIM. (The curves represent the averages of three independent experiments.)

Figure S10. Overhanging primers (**A** and **B**) used for linearization of the plasmid and recombination into the genome. The forward primer contained a 100-bp overhanging segment homologous to *rrnD* downstream from the site of insertion (brown). The reverse primer contained a 50-bp overhanging region homologous to *rrnD* upstream from the site of insertion (orange). The CRISPR-Cas9 targeting sequence (**C**) was cloned into pCRISPR plasmid (Addgene #42875) and used as crRNA targeting the Cas9 protein (expressed from Addgene #42876) to the ampicillin resistance gene of pSG76A.

Table S1. Growth parameters of BL21(DE3) (marked BL), SixPack, and Rosetta2(DE3)pLysS (marked R) strains grown in LB and AIM.

Table S2. Size and rare codon composition of the ORFs of test proteins. SixPack showed more effective protein production than the parental strain BL21(DE3) when the ratio of rare codons was about 8% or more (labeled with circles). The efficient translation of EFT1 by SixPack might be due to the extremely high ratio of one rare codon species, AGA (labeled with rectangle).

Table S3. List of 16 protein-expressing plasmid (pETDuet-1) constructs and the cloning sites used.

Table S4. Primers used for qRT-PCR. *argX*, *glyT*, *leuW*, *proL*, *argU,* and *ileX* correspond to rare; *alaU*, *gltW* and *proK* correspond to abundant (control) tRNA species.

